# Enhancing the antibacterial function of probiotic *Escherichia coli* Nissle: when less is more

**DOI:** 10.1101/2023.06.09.544192

**Authors:** Emma Bartram, Masanori Asai, Philippe Gabant, Sivaramesh Wigneshweraraj

## Abstract

Probiotic bacteria confer multiple health benefits, including preventing the growth, colonisation, or carriage of harmful bacteria in the gut. Bacteriocins are antibacterial peptides produced by diverse bacteria and their production is tightly regulated and coordinated at the transcriptional level. A popular strategy for enhancing the antibacterial properties of probiotic bacteria is to retrofit them with the ability to overproduce heterologous bacteriocins. This is often achieved from non-native constitutive promoters or in response to host or pathogen signal from synthetic promoters. How the dysregulated overproduction of heterologous bacteriocins affects the fitness and antibacterial efficacy of the retrofitted probiotic bacteria is often overlooked. We have conferred the prototypical probiotic *Escherichia coli* strain Nissle (EcN) the ability to produce McC from the wild-type promoter and two mutant promoters that allow, relative to the wild-type promoter, high and low amounts of McC production. This was done by introducing specific changes to the sequence of the wild-type promoter driving transcription of the McC operon, whilst ensuring that the modified promoters respond to native regulation. By studying the transcriptomic responses and antibacterial efficacy of the retrofitted EcN bacteria in a *Galleria mellonella* infection model of enterohemorrhagic *E. coli*, we show that EcN bacteria that produce the lowest amount of McC display the highest antibacterial efficacy with little to none undesired collateral impact on their fitness. The results highlight considerations researchers may take into account when retrofitting probiotic bacteria with heterogenous gene products for therapeutic, prophylactic or diagnostic applications.

**IMPORTRANCE:** Bacteria that resist killing by antibiotics are a major risk to modern medicine. The use of beneficial ‘probiotic’ bacteria as chassis to make antibiotic-like compounds at the site of infection in the body is emerging as a popular alternative to the use of conventional antibiotics. A potential drawback of engineering probiotic bacteria in this way is that producing antibiotic-like compounds could impart undesired side-effects on the performance of such bacteria and thereby compromise their intended use. This study highlights considerations researchers may take into account when engineering probiotic bacteria for therapeutic, prophylactic or diagnostic applications.

---

The antibiotic resistance crisis necessitates innovative approaches to manage bacterial infections. Probiotics, which are live microorganisms, have the potential to be used as prophylactic or therapeutic alternatives to antibiotics (1, 2). Probiotics can protect against enteric bacterial pathogens through various mechanisms such as competition for nutrients, activation of host defences, strengthening of the gut epithelial barrier and through the production of antibacterial peptides, including bacteriocins (2–5). Bacteriocins exhibit high structural and functional diversity (6), but their target range is often narrow and confined to bacteria closely related to the producing strain. Typically, the genes responsible for bacteriocin production are found alongside one or more cognate immunity determinants in chromosomally encoded or plasmid borne gene clusters (6). Although the production of bacteriocins confers a competitive advantage to the producing strain, it is well recognised that bacteriocin production also imparts a fitness cost on the producing strain (7–11).Therefore, the bacteriocin synthesis genes are often under tight transcriptional control and only become activated in response to metabolic or quorum dependent signals, or as part of the SOS-response (12).

In recent years, various efforts have been made to enhance the inherent antibacterial properties of probiotic bacteria by equipping them with the ability to produce heterologous bacteriocins against a pathogen of interest (13–17). *Escherichia coli* Nissle 1917 (EcN), a medically licenced probiotic, used for the treatment of gastrointestinal conditions including ulcerative colitis and infant diarrhoea, is often used as the model chassis to introduce heterologous bacteriocin genes (18–20). EcN is non-pathogenic, non-invasive, capable of colonising the intestinal tract for long periods and can provide protection against various enteropathogens by modulating host immunity, including by supressing Shiga toxin production from enterohaemorrhagic bacteria and competing directly with pathogens for nutrients (21–23). EcN naturally produces two bacteriocins, microcin H47 and microcin M. The production of microcins H47 and M is dependent on the iron availability (24). This can limit the bacteriocin-dependent antibacterial properties of EcN as iron levels can substantially fluctuate in people depending on their diet, genetic factors and intestinal health status (25). Therefore, conferring EcN (and other probiotic bacteria) with the ability to produce heterologous bacteriocins is an emerging and popular strategy to enhance the antibacterial prowess of probiotic bacteria. A survey of the current literature at the time of writing showed that, in all except one documented case, bacteriocin production was decoupled from its native regulation to achieve high amounts of the bacteriocin of interest (Fig. 1A). This was done either by introducing a synthetic promoter to constitutively drive transcription of bacteriocin genes or allow the transcription of bacteriocin genes in response to a host or pathogen signal (9, 13–16, 26). However, it is unclear whether the high amounts of bacteriocin production, decoupled from native regulation, is warranted to obtain the desired antibacterial effect. This is important to know because sustained high amounts of production of heterologous bacteriocins could impose a metabolic burden and thereby compromise the inherent beneficial traits of EcN bacteria or introduce undesired collateral physiological traits and/or fitness cost that might compromise the intended enhanced antibacterial efficacy of the engineered EcN bacteria.

**Fig. 1.**
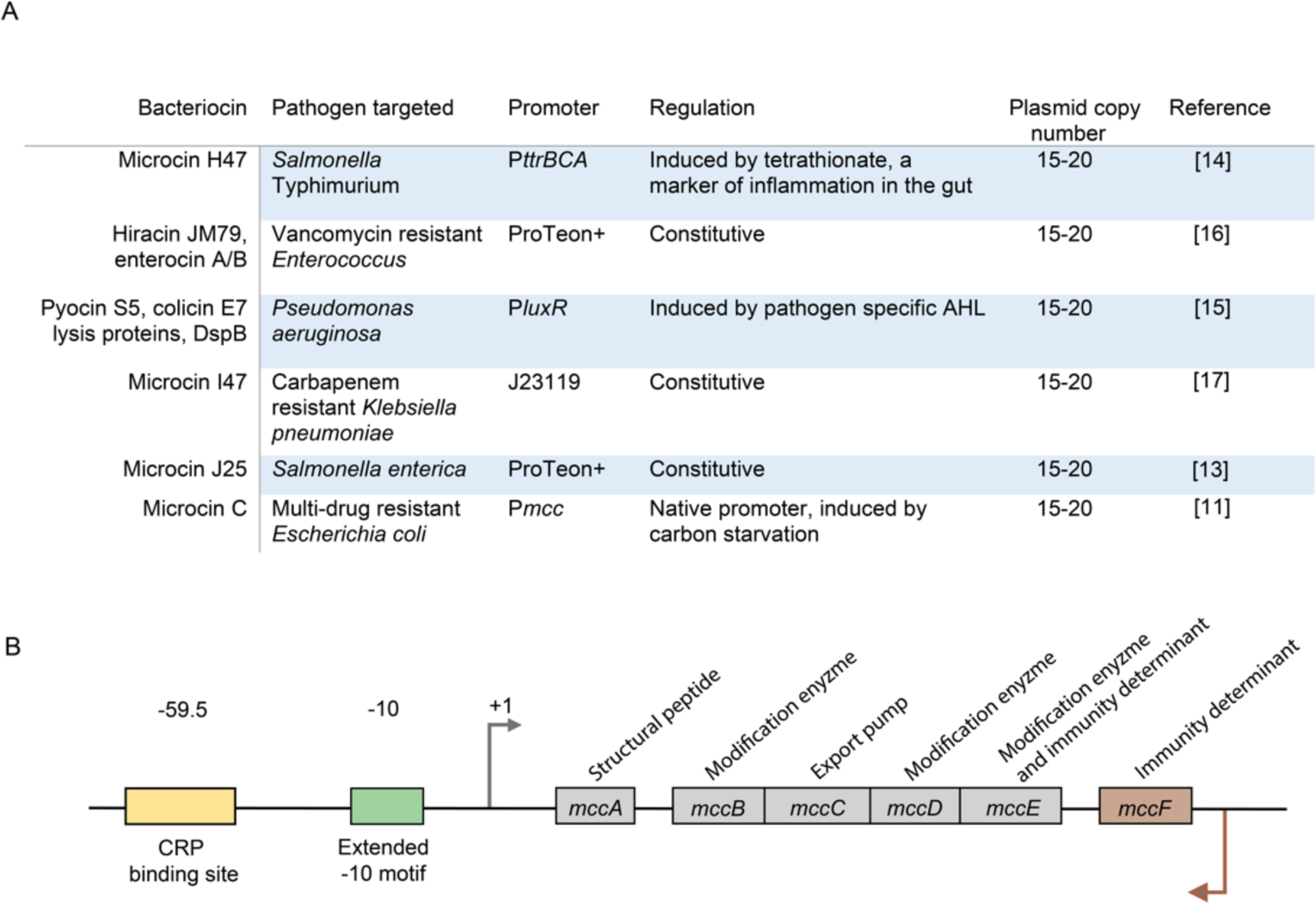
Schematic representation of the McC operon where the organisation of the structural genes (*mccA-E*) and the immunity determinant (*mccF*) are shown with respect to the transcriptional regulatory elements.

Microcin C (McC) is a narrow spectrum yet potent translational inhibitor, which is produced naturally by some *E. coli* strains and predominantly affects closely related bacteria including pathogenic *Escherichia*, *Klebsiella* and *Shigella* species (27). The genes responsible for the synthesis and secretion of, and immunity to, McC, *mccA-F* are carried on a low copy number (1-5 copies per cell) plasmid. The *mccA-E* genes are transcribed from a single operon and encode the structural peptide (MccA), enzymes (MccB, MccD and MccE), which post-translationally modify MccA to produce mature McC, the McC export pump (MccC) and an immunity determinant (MccE) (27–29) (Fig. 1B). An additional immunity determinant, MccF, is a separate transcription unit that is located downstream of *mccE* and is positioned in the opposite orientation to the *mccA-E* operon (30) (Fig. 1B). The transcription of the *mccA-E* operon is driven from an extended −10 promoter (P*mcc*; Fig. 1B). P*mcc* lacks an obvious −35 consensus element. Instead, a binding site for the catabolite repressor protein CRP is found centred ∼60 nucleotides upstream of the transcription start site of *mccA* (Fig. 1B). Therefore, optimal transcription of *mccA-E* operon is subjected to stringent regulation and relies on the CRP and the RNA polymerase (RNAP) promoter specificity factor RpoS (30–32). Hence, McC is synthesised when the optimal carbon source becomes limited (33, 34). Conversely, under nutrient replete conditions, the transcription of *mcc* genes is further repressed by the nucleoid associated proteins, HNS and Lrp (30, 32). By strategically altering the core P*mcc* sequence, we made two mutant promoters which allowed higher (P*mcc*^HIGH^) and lower (P*mcc*^LOW^) transcription of *mccA-E* genes than the WT (WT) promoter (P*mcc*^WT^). Both mutant promoters were unaltered in their dependence on CRP and RpoS for their optimal activity. The properties of EcN strains with *mccA-E* genes under P*mcc*^HIGH^ and P*mcc*^LOW^ highlight considerations that investigators might take into account when retrofitting EcN and other probiotic bacteria with bacteriocin genes or other gene products of therapeutic and prophylactic value.

## RESULTS

### The sensitivity of clinical *E. coli* isolates to McC

Our aim was to confer EcN bacteria the ability to produce different amounts of McC from P*mcc*^WT^, P*mcc*^HIGH^ and P*mcc*^LOW^ and to evaluate the consequences of doing so on the fitness and antibacterial efficacy of the EcN bacteria. Although it is well established that McC effectively stops the growth of *E. coli* and related pathogens, it is not known how sensitive clinical *E. coli* isolates are to McC and how prevalent McC resistance is. Therefore, we used a simple disc diffusion assay on Müller-Hinton agar plates to evaluate the sensitivity of a small set of clinical *E. coli* isolates (obtained from the Imperial College National Institute for Health Research, Health Protection Research Unit in Healthcare Associated Infections and Antimicrobial Resistance) to McC. The enterohemorrhagic *E. coli* O157:H7 strain EDL933 (hereafter referred to as EDL933 strain) was used as the McC sensitive reference strain. We also included broadly studied uropathogenic *E. coli* isolates CFT073 and UTI89 in our screen. None of the strains in the screen harboured any plasmids that contained the *mccA-F* genes. As McC is secreted into the culture media, McC for the disc diffusion assay was obtained by collecting and filtering the supernatant from an overnight culture of a standard laboratory *E. coli* bacteria containing a plasmid from which genes *mccA-F* were overexpressed. The size of the halo, indicating bacterial growth inhibition, surrounding the disc containing McC on plates containing the EDL933 strain, had a radius of ∼4 mm (Fig. 2). Therefore, we deemed a halo size of <2 mm as having reduced sensitivity to McC (indicated by the dotted line in Fig. 2). Results revealed that of the 21 clinical *E. coli* isolates tested, 17 were as sensitive to McC as the reference strain, including CFT073 and UTI89, under aerobic growth conditions. We noted that 2 of the clinical isolates that displayed reduced sensitivity to McC under aerobic growth conditions, showed enhanced sensitivity to McC under anaerobic growth conditions (Fig. 2). Only one clinical isolate displayed reduced sensitivity to McC under aerobic and anaerobic growth conditions. Overall, we conclude that the majority of clinical isolates of *E. coli* are sensitive to McC and the prevalence of *mccF* independent McC resistance is low.

**Fig. 2.**
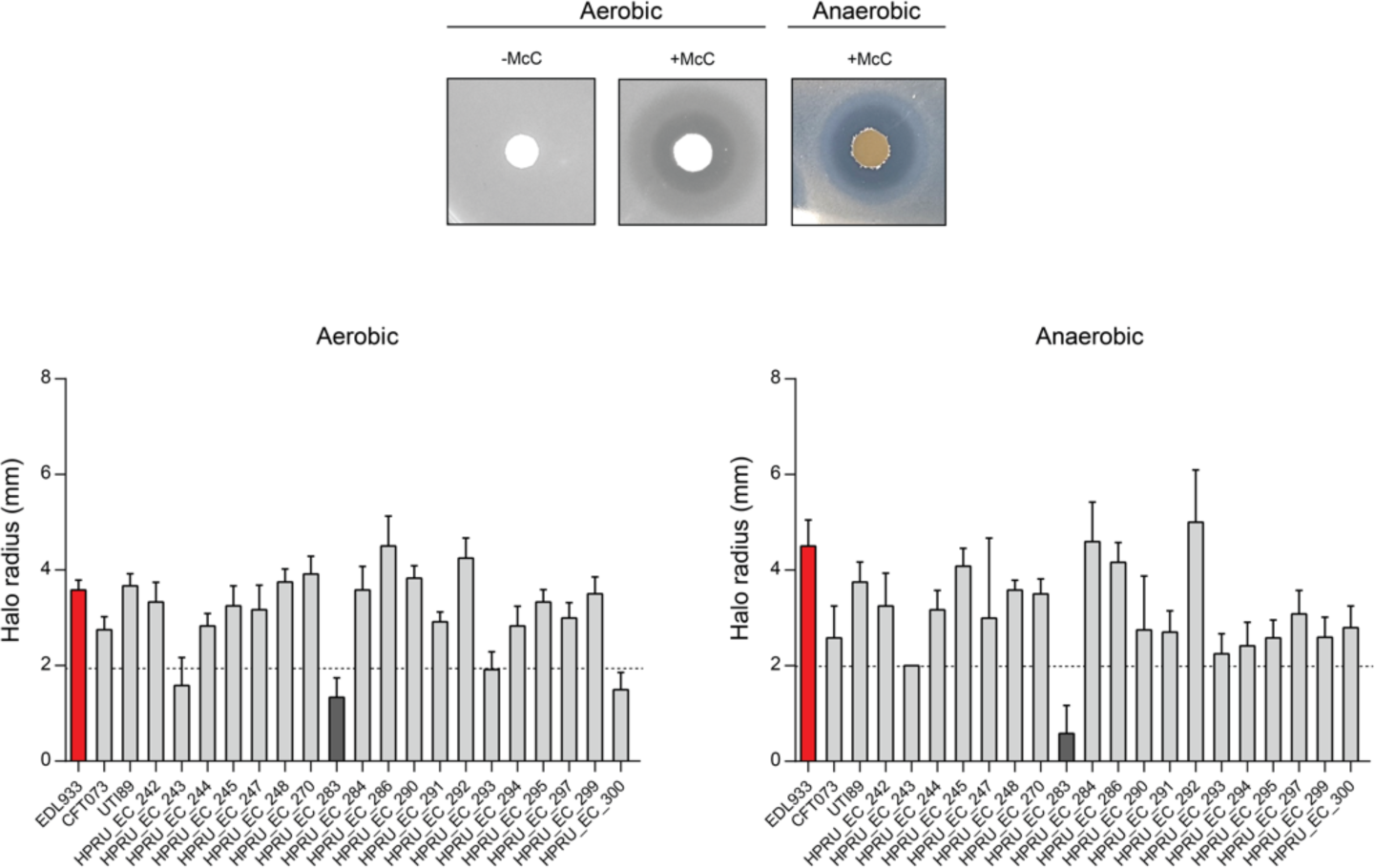
The representative images Müller-Hinton agar plates show the ‘halo’, indicating bacterial growth inhibition, surrounding the disc containing McC. The graphs show the size of the halo produced (radius, in mm) on Müller-Hinton agar plates containing the clinical *E. coli* isolates. The red bar indicates the McC sensitive reference strain *E. coli* O157:H7 strain EDL933; the dark bars indicate the clinical isolates that displayed reduced sensitivity to McC.

### P*mcc* exist in different evolutionary clades but do not display differences in activity and regulation

Prior to constructing plasmids with *mccA-E* genes under P*mcc*^WT^, P*mcc*^HIGH^ and P*mcc*^LOW^, we wanted to know how conserved the P*mcc* core promoter and upstream regulatory sequences are. Therefore, we used the nucleotide sequence of the *mccB* gene as the query to identify all *E. coli* strains available in the NCBI RefSeq genomes database that potentially contained a McC producing plasmid. We identified 185 unique *E. coli* strains from which we then compared the nucleotide sequence ∼400 base pairs upstream of the first codon of MccA. Phylogenetic analysis revealed that P*mcc* and the upstream regulatory sequences from the 185 *E. coli* strains fall into three distinct clades, with 69 and 108 of them represented in clades 1 and 2 and only 8 represented in clade 3 (Fig. 3A). The core P*mcc* sequence and the CRP binding site was conserved in all three clades (Fig. 3B). Notably, the sequences in clade 1 differed from those in clades 2 and 3 by the absence of a 21 base pair long sequence, located ∼225 nucleotides upstream of the first codon of MccA. As the majority of P*mcc* sequences either fell into clades 1 or 2, we focused our analysis on these two clades. We first wanted to determine whether the 21 base pair long insertion and other differences in the nucleotide sequence upstream of the core P*mcc* sequence affected the activity and regulation of the promoter that belong to clades 1 and 2. To do so, we fused the ∼400 base pair long sequence upstream of the first codon of MccA from a representative clade 1 and clade 2 promoter to green fluorescent protein (GFP) and placed this P*mcc*-GFP transcriptional fusion in the low copy number (10-12 per cell) plasmid pACYC to generate plasmids pEB-C1-GFP and pEB-C2-GFP (Table 1). We then measured GFP production from pEB-C1-GFP and pEB-C2-GFP initially in *E. coli* strain MG1655 grown in lysogeny broth (LB). As shown in Fig. 3C, we did not observe any difference in GFP production from pEB-C1-GFP and pEB-C2-GFP in both strains grown as batch cultures. Although bacteria containing pEB-C1-GFP and pEB-C2-GFP did not differ in GFP production at the whole population level, we considered whether they differed at the single cell level. Hence, we took bacteria after 4 h of growth (when GFP production became detectable; Fig. 3C) in LB and enumerated GFP producing individual cells by fluorescence microscopy. As shown in Fig. 3D, we did not observe any differences in the number of individual GFP fluorescent bacteria in samples containing pEB-C1-GFP and pEB-C2-GFP. As P*mcc* is dependent on RpoS and CRP for optimal activation, we next determined whether clade 1 and clade 2 promoters differ in their dependency on RpoS and ability to respond to regulation by CRP. Hence, we compared GFP production from pEB-C1-GFP and pEB-C2-GFP in a Δ*rpoS* MG1655 strain and WT MG1655 strain grown in LB supplemented with extra glucose (note that the presence of excess glucose will abrogate CRP binding to DNA), respectively. As shown in Fig. 3E, we observed a ∼2.7-fold reduction in GFP production from both pEB-C1-GFP and pEB-C2-GFP in the Δ*rpoS* MG1655 strain compared to WT MG1655 strain following 10 h of growth. This confirms previous work that showed that whilst the P*mcc* is categorised as a RpoS dependent promoter, it can be potentially used by RpoD, the major housekeeping promoter specificity factor *in E. coli*, when RpoS is unavailable (35). However, GFP production from pEB-C1-GFP and pEB-C2-GFP was fully attenuated when WT MG1655 bacteria containing pEB-C1-GFP and pEB-C2-GFP were grown in LB with extra glucose (Fig. 3F). This suggests that whilst P*mcc* might not be fully reliant on RpoS, it has a strict requirement for CRP for activation of transcription. Overall, we conclude that, despite major evolutionary differences in the upstream regulatory sequences, clade 1 and clade 2 P*mcc* promoters do not differ in their activity and regulation.

**Fig. 3.**
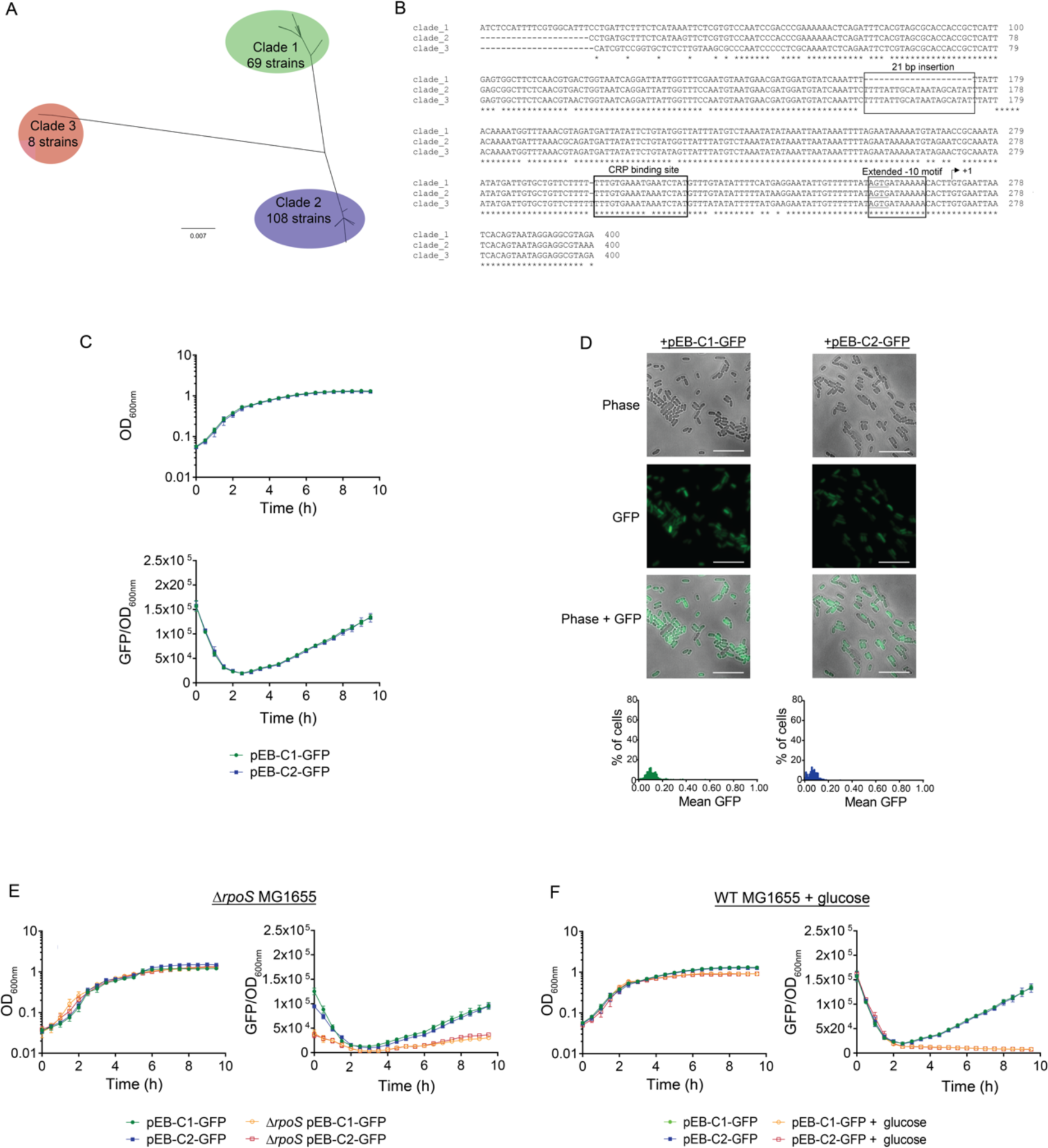
**(A)** Phylogenetic tree of P*mcc* and upstream regulatory sequence. **(B)** Sequence alignment of representative sequences from clades 1-3. The transcription start site of Pmcc is indicated and the 21 bp insertion, CRP binding site and the extended −10 motif are boxed. **(C)** Graphs showing growth (OD_600nm_) and GFP fluorescence (GFP/ OD_600nm_) from pEB-C1-GFP and pEB-C2-GFP in WT MG1655 bacteria. **(D)** Representative microscopy images of WT MG1665 bacteria containing pEB-C1-GFP or pEB-C2-GFP following 4 h of growth in LB. **(E)** Graphs showing growth (OD_600nm_) and GFP fluorescence (GFP/ OD_600nm_) from pEB-C1-GFP and pEB-C2-GFP in Δ*rpoS* MG1655 bacteria. **(F)** Graphs showing growth (OD_600nm_) and GFP fluorescence (GFP/ OD_600nm_) from pEB-C1-GFP and pEB-C2-GFP in WT MG1655 bacteria grown in LB supplemented with glucose.

### Construction and characterisation of P*mcc*^HIGH^ and P*mcc*^LOW^

Having established that clade 1 and clade 2 P*mcc* promoters do not differ in their activity and regulation, we used a representative clade 1 P*mcc* promoter to construct P*mcc*^WT^, P*mcc*^HIGH^ and P*mcc*^LOW^. To make P*mcc*^HIGH^ we introduced a suboptimal −35 sequence (5’-GTGCCA-3’ instead of the optimal 5’- TTGACA-3’ sequence) in P*mcc*^WT^ (Fig. 4A). We expected that a suboptimal −35 consensus sequence together with the extended −10 element would subtly increase RNAP binding, whilst retaining dependency on CRP, and thereby lead to an overall increased transcriptional output from the modified promoter. Conversely, to make P*mcc*^LOW^, we swapped the first 5’-TG-3’ of the TGTGn extended −10 motif of P*mcc*^WT^ to a 5’-GT-3’ (Fig. 4A). We then fused P*mcc*^WT^, P*mcc*^HIGH^ and P*mcc*^LOW^ to GFP in pACYC to generate pEB-P*mcc*^WT^-GFP, pEB-P*mcc*^HIGH^-GFP and pEB-P*mcc*^LOW^-GFP (Table 1) and measured promoter activity in *E. coli* MG1655 and EcN bacteria. As shown in Fig. 4B, as expected, in *E. coli* MG1655, the maximal rate of GFP accumulation from P*mcc*^HIGH^ was ∼1.4 fold higher than the rate of GFP accumulation from P*mcc*^WT^ (10170 GFP/OD_600nm_/hour for WT; 14633 GFP/OD_600nm_/hour for high). This meant that after 10 h of growth in LB, the amount of GFP made from P*mcc*^HIGH^ exceeded that made from P*mcc*^WT^ by ∼1.6-fold. Conversely, the rate of GFP accumulation from P*mcc*^LOW^ was ∼2.9-fold lower than the rate of GFP accumulation from P*mcc*^WT^ (3548 GFP/OD_600nm_/hour for low). This meant that after 10 h of growth in LB, the amount of GFP made from P*mcc*^LOW^ was reduced by ∼3.4-fold compared to that made from P*mcc*^WT^. Similarly, in EcN bacteria, the rate of GFP accumulation from P*mcc*^HIGH^ was ∼1.8-fold higher than the rate of GFP accumulation from P*mcc*^WT^ (2679 GFP/OD_600nm_/hour for WT; 4900 GFP/OD_600nm_/hour for high). This meant that after 10 h of growth in LB, the amount of GFP made from P*mcc*^HIGH^ exceeded that made from P*mcc*^WT^ by ∼1.9-fold. Conversely, the rate of GFP accumulation from P*mcc*^LOW^ was ∼2.0-fold lower than the rate of GFP accumulation from P*mcc*^WT^ (1357 GFP/OD_600nm_/hour for low). This meant that after 10 h of growth in LB, the amount of GFP made from P*mcc*^LOW^ was reduced by 2.3-fold compared to that made from P*mcc*^WT^. In both, *E. coli* MG1655 and EcN bacteria, the amount of GFP produced from P*mcc*^HIGH^ and P*mcc*^LOW^ differed by ∼5.4 and ∼4.5-fold, respectively. As shown in Fig. 4D, and expected (see above), in the Δ*rpoS* MG1655 strain, the activity of all three promoters was reduced by ∼1.5-2-fold compared to their relative activity in WT MG1655 strain. However, GFP production was fully abolished when WT MG1655 bacteria containing P*mcc*^WT^, P*mcc*^HIGH^ and P*mcc*^LOW^ were grown in LB with extra glucose (Fig. 4E). Overall, we conclude that, P*mcc*^HIGH^ and P*mcc*^LOW^ display the expected difference in transcription activity compared to P*mcc*^WT^ and that the changes made to P*mcc*^WT^ sequence to generate P*mcc*^HIGH^ and P*mcc*^LOW^ do not affect their regulation.

**Fig. 4.**
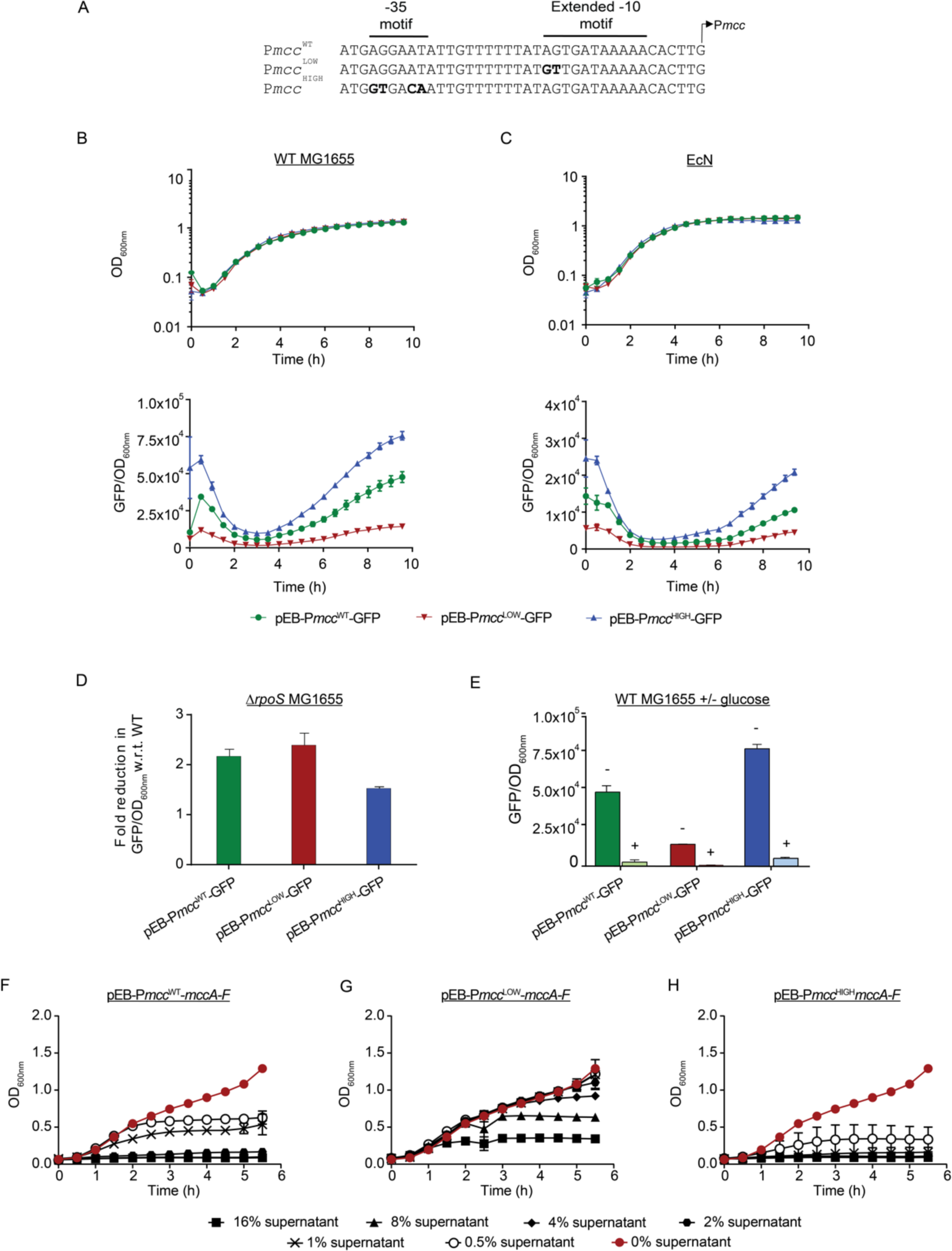
**(A)** Sequences showing the nucleotide changes introduced (in bold) into P*mcc*^WT^ to generate P*mcc*^HIGH^ and P*mcc*^LOW^. **(B)** Graphs showing growth (OD_600nm_) and GFP fluorescence (GFP/ OD_600nm_) from pEB-P*mcc*^WT^-GFP, pEB-P*mcc*^HIGH^-GFP and pEB-P*mcc*^LOW^-GFP in the *E. coli* MG1655 strain. **(C)** As in (B) but in the EcN strain. **(D)** Bar chart showing GFP fluorescence (GFP/ OD_600nm_) following 10 h of growth in in *E. coli* Δ*rpoS* MG1655 strain. **(E)** As in (D) but in the *E. coli* MG1655 strain in the absence and presence of glucose. **(F)-(H)** Graphs showing inhibition of growth of *E. coli* EDL933 in the presence of different amounts of McC containing supernatant from EcN bacteria containing pEB-P*mcc*^WT^-*mccA-F* (F), pEB-P*mcc*^HIGH^-*mccA-F* (G) and pEB-P*mcc*^LOW^-*mccA-F* (H).

Next, we replaced the GFP in pEB-P*mcc*^WT^-GFP, pEB-P*mcc*^HIGH^-GFP and pEB-P*mcc*^LOW^-GFP with *mccA-F* gene cassette (as shown in Fig. 1B) to generate pEB-P*mcc*^WT^-*mccA-F*, pEB-P*mcc*^HIGH^-*mccA-F* and pEB-P*mcc*^LOW^-*mccA-F* (Table 1) and determined whether the differences in GFP activity from the three promoters corresponded to McC production in EcN. To do so, EcN bacteria harbouring pEB-P*mcc*^WT^-*mccA-F*, pEB-P*mcc*^HIGH^-*mccA-F* and pEB-P*mcc*^LOW^-*mccA-F* were grown overnight in LB, their supernatant harvested, filtered, and used as the source of McC. Different amounts (0-16% (v/v)) of this supernatant was then used to supplement fresh LB media in 96-well plates, which were inoculated with enterohemorrhagic *E. coli* EDL933. As shown in Fig. 4F, ≥2% (v/v) of supernatant from EcN with pEB-P*mcc*^WT^ resulted in complete growth attenuation of *E. coli* EDL933; and moderate growth was only detected with ∼1% (v/v) of supernatant from EcN with pEB-P*mcc*^WT^-*mccA-F*. In contrast, in experiments with supernatant from EcN with pEB-P*mcc*^LOW^-*mccA-F*, growth of *E. coli* EDL933 comparable to that in wells with 0% (v/v) supernatant was detected with up to ∼4% (v/v) supernatant (Fig. 4G). However, in experiments with supernatant from EcN with pEB-P*mcc*^HIGH^-*mccA-F*, only ∼0.5% (v/v) of supernatant supported some growth of *E. coli* EDL933 and no growth was detected in well with ≥0.5% (v/v) supernatant (Fig. 4H). We estimate that ∼2-fold more McC is produced by EcN containing pEB-P*mcc*^HIGH^-*mccA-F* than in EcN containing pEB-P*mcc*^WT^, and, conversely, >8-fold less McC is produced by EcN containing pEB-P*mcc*^LOW^-*mccA-F* than in EcN containing pEB-P*mcc*^WT^. Hence, the McC produced by EcN containing pEB-P*mcc*^HIGH^-*mccA-F* and EcN containing pEB-P*mcc*^LOW^-*mccA-F* differs by >16-fold. Overall, we conclude that P*mcc*^HIGH^ and P*mcc*^LOW^ confer EcN the ability to make high and low amounts, respectively, of McC compared to from P*mcc*^WT^.

### The impact of producing different amounts of McC on EcN fitness

To determine how producing high and low amounts of McC from pEB-P*mcc*^HIGH^-*mccA-F* and pEB-P*mcc*^LOW^-*mccA-F* affects the growth dynamics and long-term viability of EcN, we conducted growth and viability assays in LB. An additional control included EcN bacteria containing pEB-P*mcc*^MUT^-*mccA-F* (in which the core P*mcc* sequence was scrambled to inactivate the promoter; Fig. S1). As shown in Fig. 5A, the growth dynamics of EcN bacteria containing pEB-P*mcc*^WT^-*mccA-F*, pEB-P*mcc*^HIGH^-*mccA-F* and pEB-P*mcc*^LOW^-*mccA-F* did not markedly differ during exponential phase of growth. Nonetheless, we detect a moderate adverse effect on growth when the bacteria entered the stationary phase of growth (which is when McC production starts from P*mcc* (see Fig. 4B) and the adversity on growth correlated with the amount of McC produced (Fig. 5A). Put simply, it appears that the amount of McC produced inversely correlates with the ability of the EcN to accumulate biomass in LB over a 24 h period of time. Thus, the production of lower amounts of McC in EcN bacteria containing pEB-P*mcc*^LOW^-*mccA-F* did not seem to affect the ability of EcN to accumulate biomass (Fig. 5A, compare pEB-P*mcc*^LOW^-*mccA-F* with pEB-P*mcc*^MUT^-*mccA-F*).

**Fig. 5.**
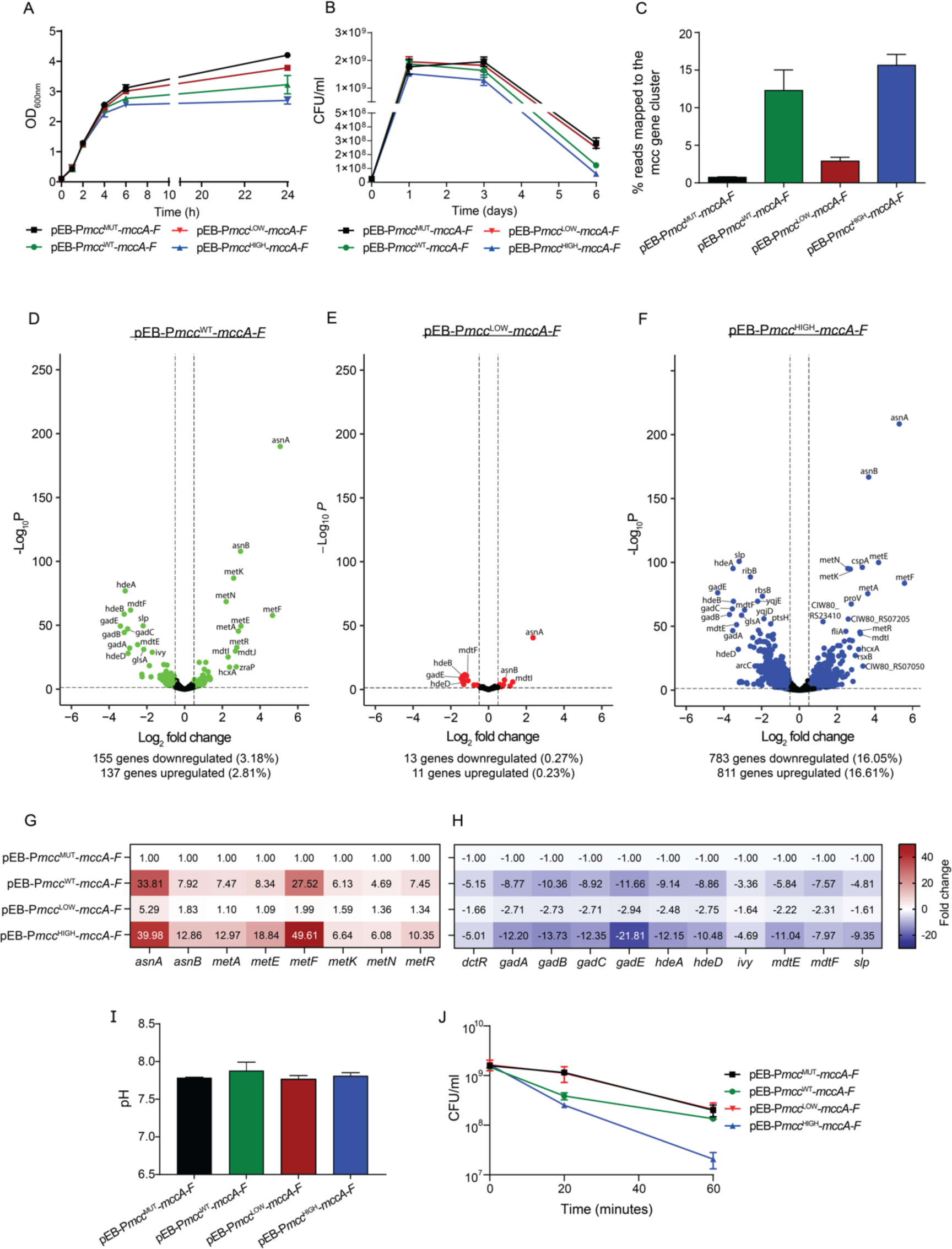
**(A)** Graph showing the growth dynamics of EcN bacteria containing pEB-P*mcc*^WT^-*mccA-F*, pEB-P*mcc*^HIGH^-*mccA-F*, pEB-P*mcc*^LOW^-*mccA-F* and pEB-P*mcc*^MUT^-*mccA-F* in LB. **(B)** Graph showing the viability over time of EcN bacteria containing pEB-P*mcc*^WT^-*mccA-F*, pEB-P*mcc*^HIGH^-*mccA-F*, pEB-P*mcc*^LOW^-*mccA-F* and pEB-P*mcc*^MUT^-*mccA-F* in LB. **(C)** Graph showing the number of reads as a percentage of total reads in the RNA-seq data that mapped to the *mccA-F* genes in EcN bacteria containing pEB-P*mcc*^WT^-*mccA-F*, pEB-P*mcc*^HIGH^-*mccA-F*, pEB-P*mcc*^LOW^-*mccA-F* and pEB-P*mcc*^MUT^-*mccA-F*. **(D)-(F)** Volcano plots showing differentially expressed genes in EcN bacteria containing pEB-P*mcc*^WT^-*mccA-F* (D), pEB-P*mcc*^LOW^-*mccA-F* (E) and pEB-P*mcc*^HIGH^-*mccA-F* as log_2_ from EcN bacteria containing pEB-P*mcc*^MUT^-*mccA-F*. **(G)** Heatmap showing differentially upregulated genes associated with asparagine and methionine biosynthesis. **(H)** Heatmap showing differentially downregulated genes associated with conferring acid resistance. **(I)** Graph showing the pH of LB media after 6 h of growth of EcN bacteria containing pEB-P*mcc*^MUT^-*mccA-F*, pEB-P*mcc*^WT^-*mccA-F*, pEB-P*mcc*^HIGH^-*mccA-F* and pEB-P*mcc*^LOW^-*mccA-F*. **(J)** Graph showing the viability over time of EcN bacteria containing pEB-P*mcc*^WT^-*mccA-F*, pEB-P*mcc*^HIGH^-*mccA-F*, pEB-P*mcc*^LOW^-*mccA-F* and pEB-P*mcc*^MUT^-*mccA-F* in synthetic gastric juice.

To enumerate the proportion of viable cells in the population of EcN bacteria making different amounts of McC, we used overnight cultures of EcN to inoculate fresh LB media 2.5 x 10^7^ CFU and measured colony forming units (CFU) daily over a period of 6 days. As expected, on day 1, the adverse effect of McC production on growth (Fig. 5A) was reflected in the proportion of viable cells in the population of EcN bacteria producing different amounts of McC (Fig. 5B). From day 1 onwards the rate of decline in the proportion of viable cells in the population of EcN bacteria producing different amounts of McC was similar and by day 6, the proportion of viable cells in the population of EcN bacteria with pEB-P*mcc*^HIGH^-*mccA-F* was ∼4.6 fold less than the proportion of viable cells in the population of EcN bacteria containing pEB-P*mcc*^LOW^-*mccA-F* (Fig. 5B). Notably, the proportion of viable cells in the population of EcN bacteria containing pEB-P*mcc*^LOW^-*mccA-F* and pEB-P*mcc*^MUT^-*mccA-F* did not differ. Overall, EcN bacteria retrofitted with the ability to make McC from P*mcc*^LOW^ display better growth and fitness characteristics than bacteria that produce McC from P*mcc*^WT^ or P*mcc*^HIGH^.

To determine how producing elevated and reduced amounts of McC affected the fitness of EcN in more depth, we compared the transcriptomes of EcN bacteria containing pEB-P*mcc*^WT^-*mccA-F*, pEB-P*mcc*^HIGH^-*mccA-F* and pEB-P*mcc*^LOW^-*mccA-F* to that of EcN bacteria containing pEB-P*mcc*^MUT^-*mccA-F*. For the transcriptomics analysis, bacteria were grown in LB and harvested following 6 h when the cultures were in the early-stationary phase (recall that this is ∼2 h after McC production has started under our conditions). As expected, the number of transcript reads as a percentage of total transcript reads that mapped to the *mccA-F* genes in EcN bacteria containing pEB-P*mcc*^WT^-*mccA-F*, pEB-P*mcc*^HIGH^-*mccA-F*, pEB-P*mcc*^LOW^-*mccA-F* and pEB-P*mcc*^MUT^-*mccA-F* correlated with the expected amount of *mccA-F* mRNA made from the respective promoters (Fig. 5C). We note that, in the case of EcN bacteria containing pEB-P*mcc*^HIGH^-*mccA-F*, ∼15% of total RNA in the cell corresponded to the mcc transcripts. For the comparative analyses of the transcriptomes, we defined differentially expressed genes as those with expression levels changed ≥2-fold with a false discovery rate adjusted P<0.05. As shown in Fig. 5D, ∼6% of the genes in EcN containing pEB-P*mcc*^WT^-*mccA-F* was differentially expressed compared to EcN containing pEB-P*mcc*^MUT^-*mccA-F*. In contrast, ∼32% of genes in EcN containing pEB-P*mcc*^HIGH^-*mccA-F* were differentially expressed compare to EcN containing pEB-P*mcc*^MUT^-*mccA-F* (Fig. 5F). However, only ∼0.5% of genes in EcN containing pEB-P*mcc*^LOW^-*mccA-F* were differentially expressed compared to EcN containing pEB-P*mcc*^MUT^-*mccA-F* (Fig. 5E). Notably, in EcN bacteria containing pEB-P*mcc*^WT^-*mccA-F* and pEB-P*mcc*^HIGH^-*mccA-F*, some of the genes that were upregulated by 4-fold or higher were associated with asparagine (*asnA* and *asnB*) and methionine biosynthesis (*metA, metE, metF, metK and metR*) and uptake (*metN*) (Fig. 5G). This could reflect the need for high levels of asparagine, methionine and the methionine derivative S-adenosylmethionine (SAM) for McC production: Translation of the precursor heptapeptide MccA (MRTGNAN) requires two asparagine and one methionine molecules; methionine is further depleted as an indirect result of the McC maturation process, in which terminal asparagine of MccA is converted to a modified aspartate residue (36). The second stage of this process requires one molecule of SAM, which is synthesised from methionine by S-adenosylmethionine synthetase, the product of *metK* (36). Conversely, many genes that were downregulated in EcN bacteria containing pEB-P*mcc*^WT^-*mccA-F* and pEB-P*mcc*^HIGH^-*mccA-F* were associated with conferring acid resistance (*hdeA, hdeD, gadA, gadB, gadC, gadE, slp, dctR, mdtE* and *mdtF*) (Fig. 5H). Hence, we considered whether the downregulation of acid stress resistance genes was reflective of a decrease in the pH of the growth media that occurred as a consequence of McC production. However, as shown in Fig. 5I, the pH of the growth media following 6 h of growth of EcN bacteria containing pEB-P*mcc*^WT^-*mccA-F*, pEB-P*mcc*^HIGH^-*mccA-F*, pEB-P*mcc*^LOW^-*mccA-F* and pEB-P*mcc*^MUT^-*mccA-F* were similar and averaged around pH 7.75. We noted *ivy*, the product of which is an inhibitor of lysozyme, a constituent of the mammalian gut, is also downregulated in EcN bacteria containing pEB-P*mcc*^WT^-*mccA-F* and pEB-P*mcc*^HIGH^-*mccA-F* (Fig. 5H). EcN bacteria must be able to adapt to the predominantly acidic gut environment to effectively execute their intended antibacterial function. Therefore, we considered whether the downregulation of *hdeA, hdeD, gadA, gadB, gadC, gadE, slp, dctR, ivy, mdtE* and *mdtF* imparts a fitness disadvantage on EcN bacteria producing elevated amounts of McC in the gut environment. To test this, we harvested EcN bacteria following 6 h of growth in LB and inoculated them into synthetic gastric fluid media (which had a pH of ∼2.5 and contained lysozyme) and enumerated the number of viable bacteria following 20 min and 1 h of incubation. Results in Fig. 5J show that whilst ∼10% of EcN bacteria containing P*mcc*^WT^-*mccA-F*, pEB-P*mcc*^LOW^-*mccA-F* and pEB-P*mcc*^MUT^-*mccA-F* survived 1 h of exposure to synthetic gastric fluid media, >99% of EcN bacteria containing pEB-P*mcc*^HIGH^-*mccA-F* did not survive this challenge. Overall, we conclude that elevated production of McC results in metabolic perturbations that compromises the overall fitness of EcN bacteria.

### The antibacterial efficacy of engineered EcN bacteria

Having established that producing high amounts of McC imparts a fitness cost on EcN bacteria, we investigated the antibacterial efficacy of EcN bacteria containing pEB-P*mcc*^WT^-*mccA-F*, pEB-P*mcc*^HIGH^-*mccA-F* and pEB-P*mcc*^LOW^-*mccA-F*. For this, we incubated different number of cells of EcN bacteria containing pEB-P*mcc*^WT^-*mccA-F*, pEB-P*mcc*^HIGH^-*mccA-F* and pEB-P*mcc*^LOW^-*mccA-F* and *E. coli* EDL933 bacteria in LB and enumerated the CFU of *E. coli* EDL933 bacteria following ∼24 h of co-culturing. In these assays, we used *E. coli* EDL933 harbouring the kanamycin resistance gene for specific selection of *E. coli* EDL933 cells. Initially, however, we used the EcN bacteria containing pEB-P*mcc*^MUT^-*mccA-F* to evaluate the inherent, i.e., McC independent, antibacterial activity of EcN. Results in Fig. 6A show that, when ∼2 x 10^7^ EcN bacteria containing pEB-P*mcc*^MUT^-*mccA-F* and *E. coli* EDL933 were co-cultured for 24 h in LB, the was a ∼50-80% reduction in the CFU of *E. coli* EDL933 cells compared to the *E. coli* EDL933 monoculture incubated for the same period of time. Therefore, to differentiate between McC independent and dependent antibacterial activity of EcN, we expressed the reduction in CFU of *E. coli* EDL933 when co-cultured with EcN bacteria containing pEB-P*mcc*^WT^-*mccA-F*, pEB-P*mcc*^HIGH^-*mccA-F* and pEB-P*mcc*^LOW^-*mccA-F* as a percentage of CFU of *E. coli* EDL933 when co-cultured with EcN containing pEB-P*mcc*^MUT^-*mccA-F* (referred to as McC dependent antibacterial activity in Fig. 6B; see below). Notably, we did not detect any differences in the antibacterial activity of EcN containing pEB-P*mcc*^WT^-*mccA-F*, pEB-P*mcc*^HIGH^-*mccA-F* and pEB-P*mcc*^LOW^-*mccA-F* co-cultured with 1:1 (Fig. 6B) or 1:1000 (Fig. 6C) *E. coli* EDL933 cells or when the McC non-sensitive *Staphylococcus aureus* cells were co-cultured with EcN and *E. coli* EDL933 cells at 1:1:1 amounts (Fig. 6D). We conclude that, regardless of the amount of McC produced by EcN containing pEB-P*mcc*^WT^-*mccA-F*, pEB-P*mcc*^HIGH^-*mccA-F* and pEB-P*mcc*^LOW^-*mccA-F*, the McC produced is sufficient to eliminate *E. coli* EDL933 under our co-culture conditions. Therefore, to better delineate the antibacterial efficacy of EcN containing pEB-P*mcc*^WT^-*mccA-F*, pEB-P*mcc*^HIGH^-*mccA-F* and pEB-P*mcc*^LOW^-*mccA-F*, we evaluated it in the context of a *Galleria mellonella* larvae as an *in vivo* infection model for *E. coli* EDL933. Our plan was to prophylactically treat the larvae with a fixed dose of EcN bacteria containing pEB-P*mcc*^WT^-*mccA-F*, pEB-P*mcc*^HIGH^-*mccA-F* or pEB-P*mcc*^LOW^-*mccA-F* and pEB-P*mcc*^MUT^-*mccA-F* for two days, before challenging them with a lethal dose of *E. coli* EDL933 and enumerating the surviving larvae over a period of eight days. Initially, we injected the larvae with different doses of EcN bacteria containing pEB-P*mcc*^MUT^-*mccA-F*, pEB-P*mcc*^WT^-*mccA-F*, pEB-P*mcc*^HIGH^-*mccA-F* or pEB-P*mcc*^LOW^-*mccA-F* or *E. coli* EDL933 to determine the appropriate dosages of the different bacteria to use in the planned experiment. Results revealed that 1×10^4^ EcN cells is the maximum prophylactic dosage we could use to inject into the larvae without killing the larvae over a period of 8 days (Fig. 6C). Results also revealed that a dosage of 1×10^5^ *E. coli* EDL933 cells killed ∼66% of larvae just 24 h following injection (Fig. 6C). Therefore, for our planned experiment, we decided on prophylactically treating the larvae with 1×10^4^ EcN cells containing pEB-P*mcc*^MUT^-*mccA-F*, pEB-P*mcc*^WT^-*mccA-F*, pEB-P*mcc*^HIGH^-*mccA-F* or pEB-P*mcc*^LOW^-*mccA-F* and, following two days of incubation, to challenge the treated larvae with 1×10^5^ *E. coli* EDL933 cells. As shown in Fig. 6D, following 2 days after the challenge with *E. coli* EDL933, only ∼22% of untreated larvae and ∼30% larvae which were prophylactically treated with EcN bacteria containing pEB-P*mcc*^MUT^-*mccA-F* survived. The survivability of the larvae was markedly improved when the larvae were prophylactically treated with EcN bacteria producing McC. We observed that EcN bacteria with pEB-P*mcc*^LOW^-*mccA-F* is consistently better at protecting the larvae against killing by *E. coli* EDL933 compared to EcN bacteria with pEB-P*mcc*^WT^-*mccA-F* or pEB-P*mcc*^HIGH^-*mccA-F*. Overall, we conclude that, expectedly, the ability to produce McC increases the antibacterial effectiveness of EcN, but, consistent with the metabolic perturbations and associated fitness cost of producing McC, the EcN bacteria producing lower amounts of McC display markedly better antibacterial prowess than EcN bacteria producing relatively higher amounts of McC.

**Fig. 6.**
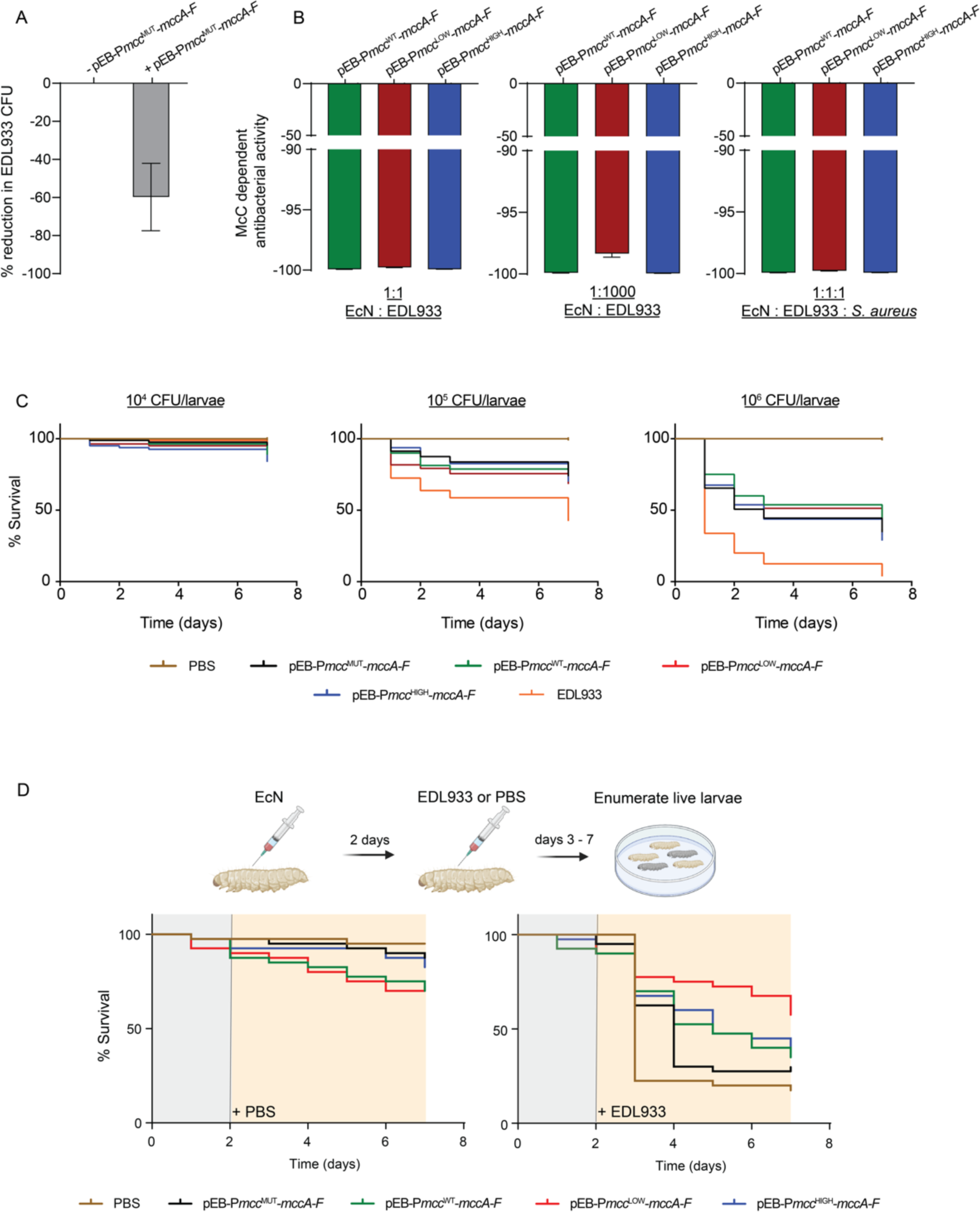
**(A)** Bar chart showing the percentage reduction in colony forming unit (CFU) when EcN bacteria containing pEB-P*mcc*^MUT^-*mccA-F* and *E. coli* EDL933 are co-cultured. **(B)** Bar chart showing the percentage reduction of *E. coli* EDL933 CFU by EcN bacteria containing pEB-P*mcc*^WT^-*mccA-F*, pEB-P*mcc*^HIGH^-*mccA-F* and pEB-P*mcc*^LOW^-*mccA-F* (see text for details). **(C)** Mortality curves of *G. mellonella* larvae infected with different amounts of EcN bacteria containing pEB-P*mcc*^WT^-*mccA-F*, pEB-P*mcc*^HIGH^-*mccA-F* and pEB-P*mcc*^LOW^-*mccA-F* or *E. coli* strain EDL933. **(D)** Mortality curves of *G. mellonella* larvae that were pre-treated with EcN bacteria pEB-P*mcc*^WT^-*mccA-F*, pEB-P*mcc*^HIGH^-*mccA-F* and pEB-P*mcc*^LOW^-*mccA-F* before infecting with *E. coli* EDL933.

## DISCUSSION

In the search for new antimicrobials, bacteriocins offer a promising alternative to conventional antibiotics. However, a major challenge with bacteriocins is their delivery in sufficient amounts to the site of infection. However, for infections of the gut, probiotic bacteria retrofitted with heterologous bacteriocin gene cassettes allow *in situ* production of sufficient amounts bacteriocins to manage enteropathogens. This is becoming an emerging alternative strategy to promote the clearance of various enteric pathogens with, so far, successful application in different animal models (15–17). As EcN is a prolific probiotic coloniser of the human gut and exhibits beneficial effects in various intestinal diseases, EcN is often used as a chassis for introducing bacteriocins to the gut environment. Although bacteriocin gene expression is tightly controlled at the transcriptional level in native producers, EcN bacteria retrofitted to produce heterologous bacteriocins are often engineered to overproduce them from constitutive promoters or in response to host or pathogen signal from synthetic promoters (Fig. 1A). However, it is well accepted that bacteriocin production imparts a fitness cost on bacteria, by draining the primary metabolite pool or because of imperfect immunity of the producer to the bacteriocin (7, 10). These costs can become more pronounced when bacteriocin production is decoupled from native regulation, since bacteriocin production continues even when it is no longer needed or unlikely to confer a competitive advantage onto the producer (37). The impetus for this study was to investigate whether the dysregulated overproduction of heterologous bacteriocins confers any adverse physiological traits on EcN which might ultimately compromise the antibacterial efficacy of the retrofitted EcN bacteria. Although our bacteriocin of choice was McC, because it is non-toxic and has been demonstrated to be capable of preventing diarrhoeal disease in weaned piglets when used to supplement feed (38), we envisage that the concepts derived from this study are applicable to wide range situations where EcN and other probiotic bacteria are engineered to produce heterologous gene products of interest for diverse applications.

We confirmed that McC is active against clinical *E. coli* isolates and showed that activity of McC is unaffected by availability of oxygen, which is severely limited in the large intestine (39), thus qualifying McC as a good candidate for use in probiotic engineering. A RpoS-dependent promoter, P*mcc*, drives the transcription of McC genes and strictly relies on the transcription activator CRP for expression (30, 32). The analysis of the core P*mcc* sequences and upstream regulatory regions revealed the presence of two major evolutionary clades and one smaller one. Notably, despite differing by the presence of a 21 bp insertion in the regulatory region upstream of P*mcc*, sequence divergences between the two larger clades did not result in any differences in transcriptional activity or regulation of representative P*mcc* from the two large clades. We consider this as indicative of the strong evolutionary pressure for retention of tight transcriptional regulation of McC production, thus further underscoring the impetus for this study that dysregulated overproduction of bacteriocins in EcN and other probiotic bacteria can potentially have adverse effects on the fitness and desired antibacterial efficacy of such bacteria.

We constructed two variants of P*mcc*, P*mcc*^HIGH^ and P*mcc*^LOW^, that allowed high and low amounts of McC production compared to the WT P*mcc* (P*mcc*^WT^), whilst full dependency on CRP, i.e., native regulation, was retained. Although P*mcc*^WT^ refers to the native sequence of P*mcc*, we note that the production of McC from P*mcc*^WT^ in our experimental setup is not necessarily reflective of McC levels in native producers. This is because we introduced the McC operon into a plasmid with a higher copy number (10–12) than the copy number (1–5) of plasmids on which the McC genes are typically found in native producers (40, 41). In fact, a previous study suggested that the production of McC from a non-native plasmid with a similar copy number to the plasmid used by us results in substantially higher amounts of McC produced than from any native plasmid (27). Nonetheless, under our conditions, the transcriptional activity and amount of McC produced from P*mcc*^HIGH^ and P*mcc*^LOW^ clearly followed the expected trend with respect to P*mcc*^WT^. Although we are yet to investigate the kinetics of RNA polymerase occupancy and activity on P*mcc*^WT^, P*mcc*^HIGH^ and P*mcc*^LOW^, the P*mcc* promoter variants we have used in this study might benefit applications where inducer-independent of accumulation of gene products of interest at different amounts are required.

The production of high amounts of McC has an adverse effect on the overall fitness of EcN bacteria. The transcriptomes of bacteria producing different amounts of McC indicate a strong correlation between the amount of McC produced and the extent of perturbation of metabolism. This could be due to the substantial allocation of cellular resources is needed to support the production of high amounts of McC. Notably, the production of McC leads to the dysregulation of genes associated with acid resistance and degradation of lysozyme – both of which are hallmark features of the gastric environment. Consequently, EcN bacteria producing high amounts of McC do not survive stimulated gastric conditions as well as the EcN bacteria producing low amounts of McC. The compromised fitness of EcN bacteria producing McC is also reflected in the *Galleria mellonella* larvae infection model of enterohaemorrhagic *E. coli* infection, in which EcN bacteria producing the lowest amount of McC show much higher antibacterial efficacy than EcN bacteria producing the highest amount of McC. It is possible that the fitness costs associated with high amounts of McC production potentially limit the ability of EcN bacteria to compete with the target pathogen. Alternatively, the metabolic perturbations associated with high amount of McC production could attenuate the other, McC independent, probiotic functions of EcN. Further, potential toxicity of McC on the *G. mellonella* larvae can also not be excluded.

Overall, this study underscores the proverbial saying ‘less is more’ should be an important consideration when retrofitting EcN and other probiotic bacteria with heterologous gene products for therapeutic, prophylactic or diagnostic applications. Thus, the use of promoter variants that allow production of the lowest effective amount of a gene(s) of interest in response to its native regulatory signals might be a robust strategy for future design of engineered probiotics.

## MATERIALS AND METHODS

### Computational analysis of P*mcc*

A BLAST search was conducted against the RefSeq genomes database using the *mccB* nucleotide sequence as a query. The sequence 400 bp upstream of the start codon of *mccA* was manually identified for each hit. Alignment of these sequences and construction of the phylogenetic trees was performed within Geneious. Clustal Omega with the mBed algorithm was used for the initial alignment and the Geneious Tree Builder was then used to generate trees using the Tamura-Nei genetic distance model and the neighbour-joining tree build method.

### Construction of plasmids and strains

*Escherichia coli* strains and plasmids used for this study are listed in Table S1. All plasmids except pEB-P*mcc*^LOW^-GFP were constructed using Gibson assembly(42). The *E. coli* strains ECR22 and A25922R were identified as containing the *mcc* gene cluster (see above); the sequence ∼400 bp upstream of the *mccA* genes of each of these strains was used as representative for the Clade 1 and Clade 2 transcriptional regulatory regions, respectively. These sequences were inserted directly upstream of a super-folder GFP gene in the plasmid pACYC184 (43) which was modified to remove the tetracycline resistance cassette. For promoter modification, pEB-C1-GFP was renamed pEB-P*mcc*^WT^-GFP and used as a vector for constructing pEB-P*mcc*^MUT^-GFP, pEB-P*mcc*^LOW^-GFP and pEB-P*mcc*^HIGH^-GFP. 120 bp complementary oligonucleotides encompassing the P*mcc*^HIGH^ and Pmcc^LOW^ promoter regions were commercially synthesised and cloned into pEB-P*mcc*^WT^-GFP, replacing the corresponding sequence in pEB-P*mcc*^WT^-GFP. For construction of the pEB-P*mcc*-*mccA-F* set of plasmids, the *mcc* gene cluster (from the translational start codon of *mccA*) was cloned in place of GFP on the pEB-P*mcc*-GFP set of plasmids, such that *mccA* was directly downstream of the WT and modified P*mcc* promoters. The *mcc* gene cluster was amplified from pp70, kindly provided by Konstantin Severinov (Rutgers University), and includes stand-alone immunity determinant *mccF* transcribed from an independent, native promoter (Fig. 1A). The *E. coli* EDL933 strain containing the kanamycin resistance gene downstream of *hfq* was constructed as previously described (44).

### Bacterial growth

Bacteria were grown in LB at 37 ᵒC with shaking (∼180 rpm) unless otherwise stated. Strains carrying a plasmid were grown in LB supplemented with 35 µg/ml chloramphenicol (to select for pACYC184 derivatives – see above). Overnight cultures of Δ*rpoS* MG1655 strain and EDL933 strain (if containing the kanamycin resistance gene) were supplemented with 50 µg/ml kanamycin.

### McC susceptibility disc assays

For assays done under aerobic conditions, bacteria were grown to mid-exponential phase (OD_600nm_ of ∼0.4). Two hundred microlitres of culture was mixed with 3 ml of cooled molten soft Mueller Hinton agar and poured onto a Mueller Hinton hard agar plate with a depth of 4 mm. Discs infused with 3 µl of filtered supernatant from an McC producer strain (supplied by Syngulon) were applied. The plates were incubated at 37 ᵒC for 18 h, after which they were imaged, and the sizes of halos measured. The McC susceptibility assays under anaerobic conditions were done as above but bacteria were grown in an anaerobic chamber and deoxygenised medium and agar were used and plates were incubated for 48 h in the anaerobic chamber before imaging. All data shown in figures is from 2 biological replicates.

### Plate reader assays

Bacterial cultures were set up in a 96-well plate and incubated in a BMG FlourStar Omega plate reader for 6-10 hours. Optical density (OD_600nm_) readings were taken at 30-minute intervals. For monitoring of GFP fluorescence over time, OD_600nm_ and OD_485nm_ readings were taken at 30-minute intervals. OD_485nm_ values were divided by OD_600nm_ values at each time point to correct for cell density. All data shown in the figures is from 3 biological replicates.

### McC susceptibility supernatant assays

Overnight cultures of EcN bacteria were centrifuged and washed in fresh LB to remove traces of any antibiotics and were then grown in LB without any antibiotics for 24 h. Subsequently, the supernatant of McC producing bacteria was harvested and passed through a 0.22 µM sterile filter. Amounts of McC within the supernatant were estimated via growth inhibition assays against the McC sensitive target EDL933 strain in a 96-well plate. Two-fold serial dilutions of supernatant were applied and growth monitored in a plate reader as described above. All data shown in the figures is from 3 biological replicates.

### Viability assays

Overnight cultures of bacteria were grown to stationary phase in a 50 ml falcon and subsequently incubated for 6 days. At each time point, serial dilutions of each culture were plated on LB agar plates and the number of CFU/ml calculated. All data shown in figures is from 3 biological replicates.

### Co-culture assays

Overnight cultures of EcN bacteria were centrifuged and washed in fresh LB to remove traces of any antibiotics. Co-cultures were set up in 50 ml falcon tubes. For co-cultures where a ratio of 1:1 EcN: EDL933, or 1:1:1 EcN:EDL933:non-susceptible competitor *S. aureus* were used. A starting dilution of ∼2 x 10^7^ CFU/ml of each strain was used. For co-cultures with a ratio of 1:100 or 1:1000 EcN: EDL933 ∼2 x 10^5^CFU/ml and ∼2 x 10^4^ CFU/ml of the EcN strains were used respectively, whilst the CFU of the EDL933 strain was maintained ∼2 x 10^7^ CFU/ml. Co-cultures were incubated for 24 hours, then they were washed in PBS, serially diluted, and plated onto LB agar plates containing 50 µg/ml of kanamycin (to only select for the EDL933 strain). All data shown in the figures is from 3 biological replicates.

### RNA-sequencing experiments

Bacteria were harvested during early stationary phase (OD_600nm_ of ∼2.3-3). Three biological replicates of each strain were obtained from across two independent experiments. RNA was extracted using the PureLink RNA Mini Kit from Invitrogen. Extracted RNA was sent to the Core Unit Systems Medicine at the University of Würzburg for next-generation sequencing and alignment to the reference EcN genome assembly ASM354697v1, which includes the sequences of EcN plasmids pNissle1 and pMut2. Raw counts were analysed with the DESeq2 Bioconductor package, using the Ashr package for LogFold2 shrinkage(45, 46). All McC producing strains were compared to EcN+pEB-P*mcc*^MUT^-*mccA-F*. Statistical analyses for differential gene expression were performed using R version 4.2.2. The sequencing data are available in the ArrayExpress database at EMBL-EBI (http://www.ebi.ac.uk/arrayexpress) under accession number E-MTAB-13078.

### Galleria mellonella infection model

For initial work to determine the correct bacterial dose of the EcN and EDL933 strains, overnight cultures of the bacteria were harvested and washed twice in PBS. Bacteria were then diluted to a density of 10^6^, 10^7^ and 10^8^ CFU/ml and mixed with filter sterilised 0.2% amaranth dye (w/v in PBS) at a ratio of 2:1 to aid with accurate injection. PBS and dye without bacteria were used as a negative control. *G. mellonella* larvae were randomly distributed into 5 experimental groups and injected with 15 µl of inoculum at the lower left hindleg. The larvae were incubated for 7 days in a 37 ᵒC, 5% CO_2_ incubator and survival monitored and recorded. Data shown in FIG.s is from 5 independent experiments with a cumulative total of n=80 larvae per group. For the sequential injections, *G. mellonella* were distributed into 10 experimental groups (n=20 per group). On day 1, the larvae were injected with PBS or 10^4^ CFU/ml of one the four EcN strains. The larvae were incubated in a 37 ᵒC, 5% CO_2_ incubator for two days, after which time they were injected with either 10^6^ CFU/ml EDL933 bacteria or PBS. The larvae were monitored for a further 5 days, and survival monitored and recorded. Data shown in figures is from 2 independent experiments with a cumulative total of n=40 larvae.

### pH measurements and gastric juice challenge

For pH measurements, supernatant was harvested during early stationary phase and sterile-filtered. The pH was measured using a standard pH meter and recorded. The synthetic gastric juice medium was made as previously described by Booth and Frost, with the pH adjusted to 2.5(47). For the synthetic gastric juice challenge, bacteria were harvested during early stationary phase (OD_600nm_ of ∼2.3-3), washed in PBS and resuspended in synthetic gastric juice medium and incubated for 20 min and 1 h. Serial dilutions of the bacteria were plated on LB agar plates and the number of CFU/ml calculated. Data shown in figures is from 3 biological replicates.

## ACKNOWLEDGEMENTS

Emma Bartram was funded by a BBSRC ICASE PhD studentship. Conceptualisation, S.W.; data curation, E.B. and S.W.; formal analysis, E.B. and S.W.; methodology, E.B. and M.A.; project administration, S.W. and P.B.; resources, S.W., P.B. and M.A.; software, not applicable; supervision, S.W.; validation, S.W. and P.B.; visualization, E.B; writing – original draft, S.W.; writing – review and editing, E.B., P.B., M.A. and S.W.

## SUPPLEMENTARY FIGURE LEGENDS

**Fig. S1.**
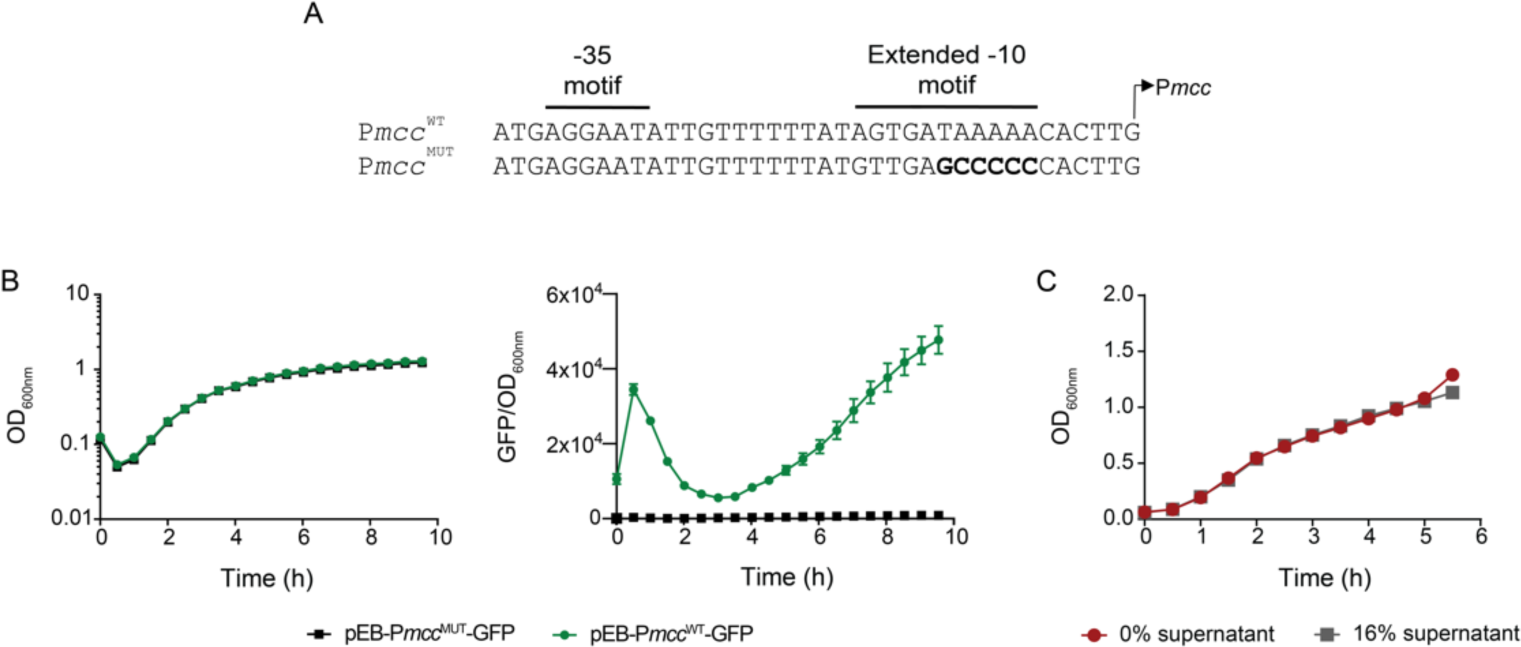
**(A)** Sequences showing the nucleotide changes introduced (in bold) into P*mcc*^WT^ to generate P*mcc*^MUT^. (B) Graphs showing growth (OD_600nm_) and GFP fluorescence (GFP/ OD_600nm_) from pEB-P*mcc*^MUT^-GFP in the *E. coli* MG1655 strain. **(C)** Graph showing growth of *E. coli* EDL933 in the presence of supernatant (16% (v/v) from EcN bacteria containing pEB-P*mcc*^MUT^-*mccA-F* (compare with Fig. 4F-FG).

**Table S1.**

| Strains |  |  |
| --- | --- | --- |
| Name | Description | Reference |
| WT MG1655 | *E. coli* K-12 F-, λ-, rph-1 | *E. coli*<br>Genetic<br>Stock Center |
| WT MG1655 + pEB-C1_GFP | WT MG1655 + pEB-C1-GFP | This study |
| WT MG1655 + pEB-C2_GFP | WT MG1655 + pEB-C2-GFP | This study |
| WT MG1655 + pEB-Pmcc<sup>WT</sup>-GFP | WT MG1655 + pEB-Pmcc<sup>WT</sup>-GFP | This study |
| WT MG1655 + pEB-Pmcc<sup>MUT</sup>-GFP | WT MG1655 + pEB-Pmcc<sup>MUT</sup>-GFP | This study |
| WT MG1655 + pEB-Pmcc<sup>LOW</sup>-GFP | WT MG1655 + pEB-Pmcc<sup>LOW</sup>-GFP | This study |
| WT MG1655 + pEB-Pmcc<sup>HIGH</sup>-GFP | WT MG1655 + pEB-Pmcc<sup>HIGH</sup>-GFP | This study |
| ΔrpoS MG1655 | MG1655 ΔrpoS::kan | |
| ΔrpoS MG1655 + pEB-C1_GFP | MG1655 ΔrpoS + pEB-C1_GFP | This study |
| ΔrpoS MG1655 + pEB-C2_GFP | MG1655 ΔrpoS + pEB-C2_GFP | This study |
| ΔrpoS MG1655 + pEB-Pmcc<sup>WT</sup>-GFP | MG1655 ΔrpoS + pEB-Pmcc<sup>WT</sup>-GFP | This study |
| ΔrpoS MG1655 + pEB-Pmcc<sup>MUT</sup>-GFP | MG1655 ΔrpoS + pEB-Pmcc<sup>MUT</sup>-GFP | This study |
| ΔrpoS MG1655 + pEB-Pmcc<sup>LOW</sup>-GFP | MG1655 ΔrpoS + pEB-Pmcc<sup>LOW</sup>-GFP | This study |
| ΔrpoS MG1655 + pEB-Pmcc<sup>HIGH</sup>-GFP | MG1655 ΔrpoS + pEB-Pmcc<sup>HIGH</sup>-GFP | This study |
| WT Nissle 1917 | Probiotic *E. coli* O6:K5:H1 strain | [1] |
| WT Nissle 1917 + pEB-Pmcc<sup>WT</sup>-GFP | WT Nissle 1917 + pEB-Pmcc<sup>WT</sup>-GFP | This study |
| WT Nissle 1917 + pEB-Pmcc<sup>MUT</sup>-GFP | WT Nissle 1917 + pEB-Pmcc<sup>MUT</sup>-GFP | This study |
| WT Nissle 1917 + pEB-Pmcc<sup>LOW</sup>-GFP | WT Nissle 1917 + pEB-Pmcc<sup>LOW</sup>-GFP | This study |
| WT Nissle E 1917 + pEB-Pmcc<sup>HIGH</sup>-GFP | WT Nissle 1917 + pEB-Pmcc<sup>HIGH</sup>-GFP | This study |
| WT Nissle 1917 + pEB-Pmcc<sup>WT</sup>-MCCA-F | WT Nissle 1917 + pEB-Pmcc<sup>WT</sup>-MCCA-F | This study |
| WT Nissle 1917 + pEB-Pmcc<sup>MUT</sup>-MCCA-F | WT Nissle 1917 + pEB-Pmcc<sup>MUT</sup>-MCCA-F | This study |
| WT Nissle 1917 + pEB-Pmcc<sup>LOW</sup>-MCCA-F | WT Nissle 1917 + pEB-Pmcc<sup>LOW</sup>-MCCA-F | This study |
| WT Nissle 1917 + pEB-Pmcc<sup>HIGH</sup>-MCCA-F | WT Nissle 1917 + pEB-Pmcc<sup>HIGH</sup>-MCCA-F | This study |
| EDL933 | EHEC O157:H7 strain | [2] |
| EDL933 (kanamycin resistant) | EDL933 *hfq*-PAmCherry-kan | This study |

Table S1
| Plasmids |  |  |
| --- | --- | --- |
| Name | Description | Reference |
| pEB-C1-GFP/<br>pEB-*Pmcc*<sup>WT</sup>-GFP | Modified pACYC backbone (-TcR) expressing GFP under the Clade 1 *Pmcc* promoter (*Pmcc*<sup>WT</sup>) | This study |
| pEB-C2-GFP | Modified pACYC backbone (-TcR) expressing GFP under the Clade 2 *Pmcc* promoter | This study |
| pEB-*Pmcc*<sup>MUT</sup>-GFP | Modified pACYC backbone (-TcR) expressing GFP under the engineered promoter *Pmcc*<sup>MUT</sup> | This study |
| pEB-*Pmcc*<sup>LOW</sup>-GFP | Modified pACYC backbone (-TcR) expressing GFP under the engineered promoter *Pmcc*<sup>LOW</sup> | This study |
| pEB-*Pmcc*<sup>HIGH</sup>-GFP | Modified pACYC backbone (-TcR) expressing GFP under the engineered promoter *Pmcc*<sup>HIGH</sup> | This study |
| pEB-*Pmcc*<sup>WT</sup>-*mccA-F* | Modified pACYC backbone (-TcR) expressing the *mcc* operon under the native Clade 1 promoter *Pmcc*<sup>WT</sup>, and immunity determinant *mccF* under the native *mccF* promoter | This study |
| pEB-*Pmcc*<sup>MUT</sup>-*mccA-F* | Modified pACYC backbone (-TcR) expressing the *mcc* operon under the engineered promoter *Pmcc*<sup>MUT</sup>, and immunity determinant *mccF* under the native *mccF* promoter | This study |
| pEB-*Pmcc*<sup>LOW</sup>-*mccA-F* | Modified pACYC backbone (-TcR) expressing the *mcc* operon under the engineered promoter *Pmcc*<sup>LOW</sup>, and immunity determinant *mccF* under the native *mccF* promoter | This study |
| pEB-*Pmcc*<sup>HIGH</sup>-*mccA-F* | Modified pACYC backbone (-TcR) expressing the *mcc* operon under the engineered promoter *Pmcc*<sup>HIGH</sup>, and immunity determinant *mccF* under the native *mccF* promoter | This study |

